# The Extent of Edgetic Perturbations in the Human Interactome Caused by Population-Specific Mutations

**DOI:** 10.1101/2023.08.08.552329

**Authors:** Hongzhu Cui, Suhas Srinivasan, Ziyang Gao, Dmitry Korkin

## Abstract

Until recently, efforts in population genetics have been focused primarily on people of European ancestry. To attenuate the bias, global population studies, such as the 1,000 Genomes Project, have revealed differences in genetic variation across ethnic groups. How much of these differences would attribute to the population-specific traits? To answer this question, the mutation data must be linked with the functional outcomes. A new “edgotype” concept has been proposed that emphasizes the interaction-specific, “edgetic”, perturbations caused by mutations in the interacting proteins. In this work, we performed a systematic *in-silico* edgetic profiling of ∼50,000 non-synonymous SNVs (nsSNVs) from 1,000 Genomes Project by leveraging our semi-supervised learning approach SNP-IN tool on a comprehensive set of over 10,000 protein interaction complexes. We interrogated functional roles of the variants and their impact on the human interactome and compared the results with the pathogenic variants disrupting PPIs in the same interactome. Our results demonstrated that a considerable number of nsSNVs from healthy populations could rewire the interactome. We also showed that the proteins enriched with the interaction-disrupting mutations were associated with diverse functions and had implications in a broad spectrum of diseases. Further analysis indicated that distinct gene edgetic profiles among major populations could shed light on the molecular mechanisms behind the population phenotypic variances. Finally, the network analysis revealed that the disease-associated modules surprisingly harbored a higher density of interaction-disrupting mutations from the healthy populations. The variation in the cumulative network damage within these modules could potentially account for the observed disparities in disease susceptibility, which are distinctly specific to certain populations. Our work demonstrates the feasibility of a large-scale *in-silico* edgetic study and reveals insights into the orchestrated play of the population-specific mutations in the human interactome.

## INTRODUCTION

Since the completion of the Human Genome Project, scientists have made remarkable advancements in high-throughput genotyping technology, in particular in the next-generation sequencing technology (NGS) [1–3]. Together with large consortium sequencing efforts, such as the International HapMap project [4] and the 1,000 Genomes Project [5], genotyping hundreds to thousands of individuals has become a common practice. The large consortium sequencing projects have revealed that many genetic variations are population-specific [5, 6]. The genetic differences among the population datasets have been critical in interpreting the phenotypic differences between populations, and they have important implications for the human health and diseases. Despite their potential significance, population-specific single nucleotide polymorphism SNPs have received limited attention and are not yet commonly considered in clinical practice.

Along with the development of NGS technologies, genetic variation databases have been developed to help scientists make sense of the vast amount of NGS data, including 1,000 Genomes Project [7] and Single Nucleotide Polymorphism Database (dbSNP) [8]. Because many genetic variants have not been previously described, the role of computational approaches for functional annotation has become increasingly important [9, 10]. There are many bioinformatics tools for the functional annotation of genetic variations. Several recent reviews [11–14] give a comprehensive survey of state-of-art variant annotation methods. Most of the tools are either sequence-based or evolutionary conservation-based, focusing on the annotation of single nucleotide variants (SNVs) because they are easier to capture and analyze [10]. If an SNV can be mapped on the experimentally determined protein structure or a corresponding accurate homology model, then one can compute some properties using the structure information, which could, in turn, improve the accuracy of predicting the functional impact of this mutation [15]. Recently, our group developed a new computational method [16], called the SNP-IN tool (non-synonymous SNP INteraction effect prediction tool) [16], to predict the effects of non-synonymous SNVs (nsSNVs) on protein-protein interactions (PPIs), given the interaction’s experimental structure or structural model. The method leverages supervised and semi-supervised learning models, including a new Random Forest self-learning protocol. The accurate and balanced performance of the SNP-IN tool allows for the proteome-scale functional annotation of non-synonymous SNVs.

At the same time, network biology has become an important approach for the *in-silico* analyses with the goal to investigate the organizing principles of intra-cellular networks and the implications of these principles for understanding diseases [17]. Besides, a complex phenotype is rarely a consequence of a single genetic mutation—it is usually the effect of genotype-environment (GxE) interaction. This complexity is further increased by the physical interactions of the genetic factors in the context of a biological network [18], with the biological networks being a natural framework for integrating different sources of data and incorporating prior knowledge [19]. Among various types of biological networks, the protein-protein interaction (PPI) network, or interactome, gains the most attention and is arguably the most widely studied network in biology [17, 20].

Recently, a concept of “edgotype” was proposed for the PPI networks [21] that described the functional outcomes of genetic variants on PPIs and the corresponding rewiring effects on the interactome. In contrast to a traditional, genotype-based, view where a genetic variation may or may not cause the loss of a protein function, the edgetic perturbation model describes a variation as interaction-specific, *i.e*., the genetic variation may or may not cause the removal or addition of a specific PPI, while other PPIs remain unperturbed. Recent studies showed that different mutations lead to different defects in proteins, and this may cause distinct perturbations in the protein-protein interaction network [22–24]. However, profiling thousands of missense mutations experimentally using interaction assays remains a costly and labor-intensive task. For the edgetic perturbation model, our SNP-IN tool can be used to carry out the *in silico* edgetic profiling.

Edgetics is a novel approach to understanding genotype-phenotype relationships in the context of the biological network. The edgotype provides the alternative, mechanistic, explanation of a mutation’s functional impact, thus dissecting many complex genotype-to-phenotype relationships [25]. An edgetic alteration can cause the removal of one or several interactions while leaving the rest intact and functional. The alteration might have a more subtle impact on the network, not necessarily resulting in a disease phenotype [26]. More importantly, the edgetic perturbation model can often easily explain confounding genetic phenomena, such as genetic heterogeneity [27, 28]. Edgetics provides us with a similar recipe to study population genetics. With the amounts of genetic variation data from different populations around the world by the global consortiums such as the 1,000 Genomes Project [7, 29] and ENCODE [30], we have a unique opportunity to move beyond the traditional population genetics and adopt a “population edgetics” approach to study genetic differences within and between populations. We also expect that the distinct edgetic profiles—rather than the mutation data alone—can better explain why there are different disease frequency patterns and susceptibilities across different populations. Ultimately, the network-based analysis of the genetic architectures may not only shed light on the biological mechanisms underlying complex phenotypes, but also yield better ways of measuring the genetic predisposition to a certain disease in healthy individuals.

In this work, we created a comprehensive catalog of population-specific edgetic effects at the whole-interactome level using population genetics data. Specifically, our recently developed SNP-IN tool was applied to 46,599 nsSNVs collected from the 1,000 Genomes Project dataset, by leveraging the structural information on PPI complexes obtained either experimentally or through modeling. We predicted that a considerable amount of healthy population specific nsSNVs could rewire PPIs. We also showed that the proteins (and the corresponding genes) enriched with the interaction-disrupting mutations obtained from the healthy populations were, in fact, associated with diverse functions and implicated in various diseases. Our analysis indicated that some gene edgetic profiles were distinct from each other across the five major populations and could help explain the population phenotypic variance. Finally, the network analysis reveals phenotype-associated modules heavily laden with the interaction-disrupting mutations. The observable discrepancy in the accumulated damage within these modules potentially indicates a variation in disease susceptibility specific to certain populations, warranting further investigation.

## RESULTS

A systematic analysis of the effects of population-specific mutations on the human interactome was conducted using the SNP-IN Tool [16] (Fig. 1). We first collected and processed population-specific mutation data from the 1,000 Genomes Project. After that, the population-specific nsSNVs were mapped onto the structures of protein-protein interaction complexes. This structural information was necessary for the SNP-IN tool’s input to predict the mutation’s effect on the corresponding protein-protein interaction (PPI). Specifically, the SNP-IN tool annotated the impact of an nsSNV on a PPI as *neutral* (no change to the PPI), *detrimental* (loss of the PPI due to the decreased binding affinity), or *beneficial* (substantially increased binding affinity of the PPI). With the functional characterization provided by the SNP-IN tool, we then evaluated interactome-wide mapping of the mutations followed by large-scale edgetic profiling of the proteins enriched with the detrimental mutations. Based on the concept of edgotype, we investigated the potential role of the human interactome’s rewiring, caused by population-specific detrimental mutations, and the resulting phenotypic variance across the populations. Finally, we leveraged the network analysis and functional module enrichment techniques to deepen our understanding of the impact of these detrimental mutations on the human interactome.

**Figure 1.**
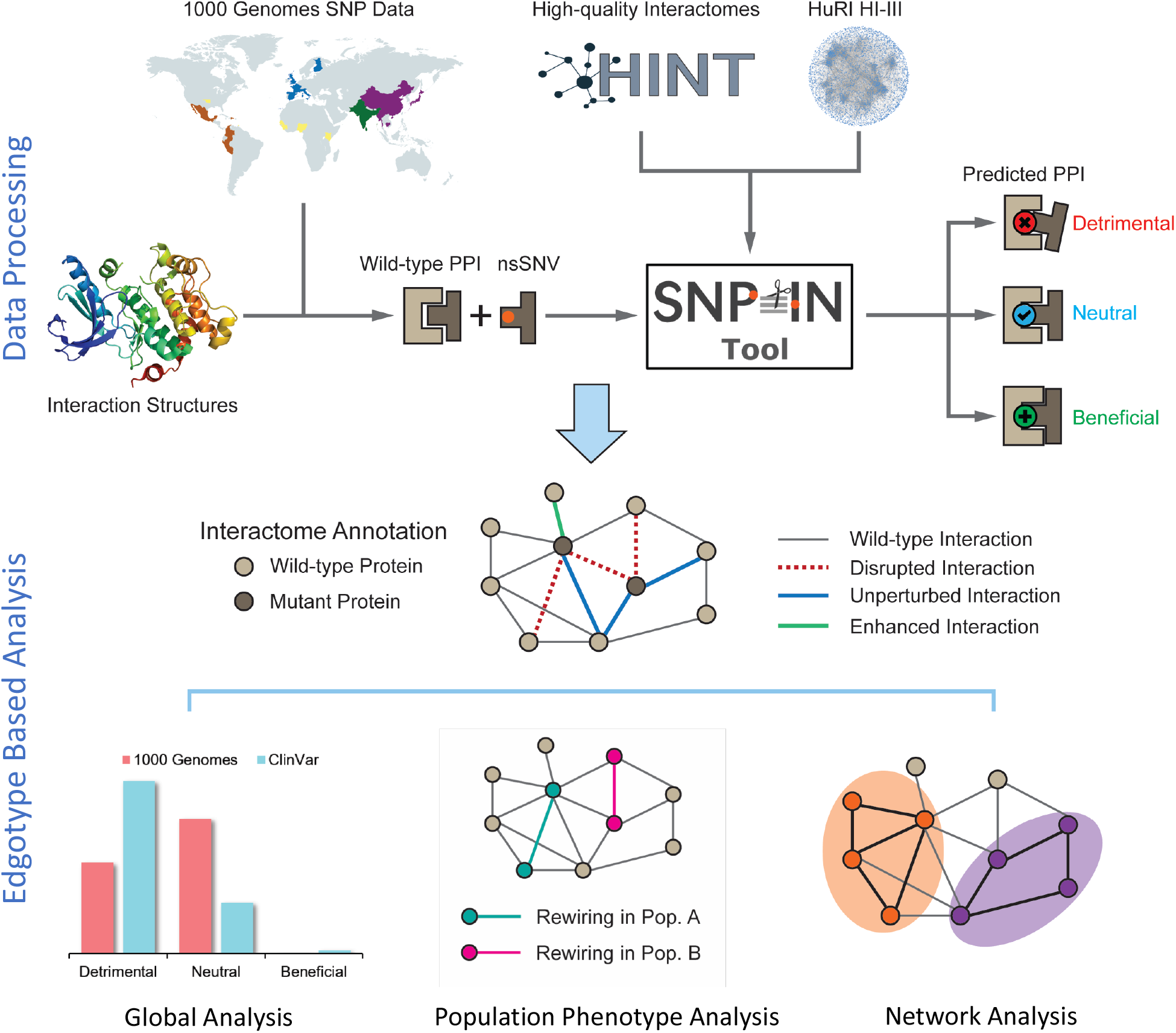
Overview of the integrated computational workflow. The workflow integrated data of three different types: population variation, protein structure, and protein interaction network. The mutation data are collected from the 1,000 Genomes Project. Then the mutations are mapped to structurally resolved protein-protein interaction complexes, and the SNP-IN tool is used for functional annotation. The analysis stage includes global evaluation of rewiring effects by mutations, edgotype-based analysis of population phenotypic variance, and network analysis of detrimental mutations.

### Interaction-disrupting nsSNVs have relatively abundant presence in healthy populations and unique evolutionary traits

Genetic variation data was extracted from the 1,000 Genomes Project dataset of more than 88 million genetic variants [5]. The majority of the variants were SNVs (84.7 million), however, the dataset also included ∼3.6 million short indels and ∼60,000 structural variants. In this work, we focused on 513,149 non-synonymous SNVs (nsSNVs), for which the corresponding residue substitution on the protein products were determined. The nsSNVs were then mapped either to experimentally obtained protein-protein interaction structures or their structural models (see Methods for more detail). Overall, 32,186 nsSNVs were mapped to 5,324 native protein-protein interaction structures, 18,119 nsSNVs to 4,258 full-length protein-protein interaction models, and 3,790 nsSNVs to 983 domain-domain interaction models (Fig. 2b). Given that a single nsSNV may be mapped to multiple experimental or computationally modeled protein-protein interaction interfaces, in total the SNP-IN tool annotated 46,599 population-specific nsSNVs based on the aforementioned mappings.

**Figure 2.**
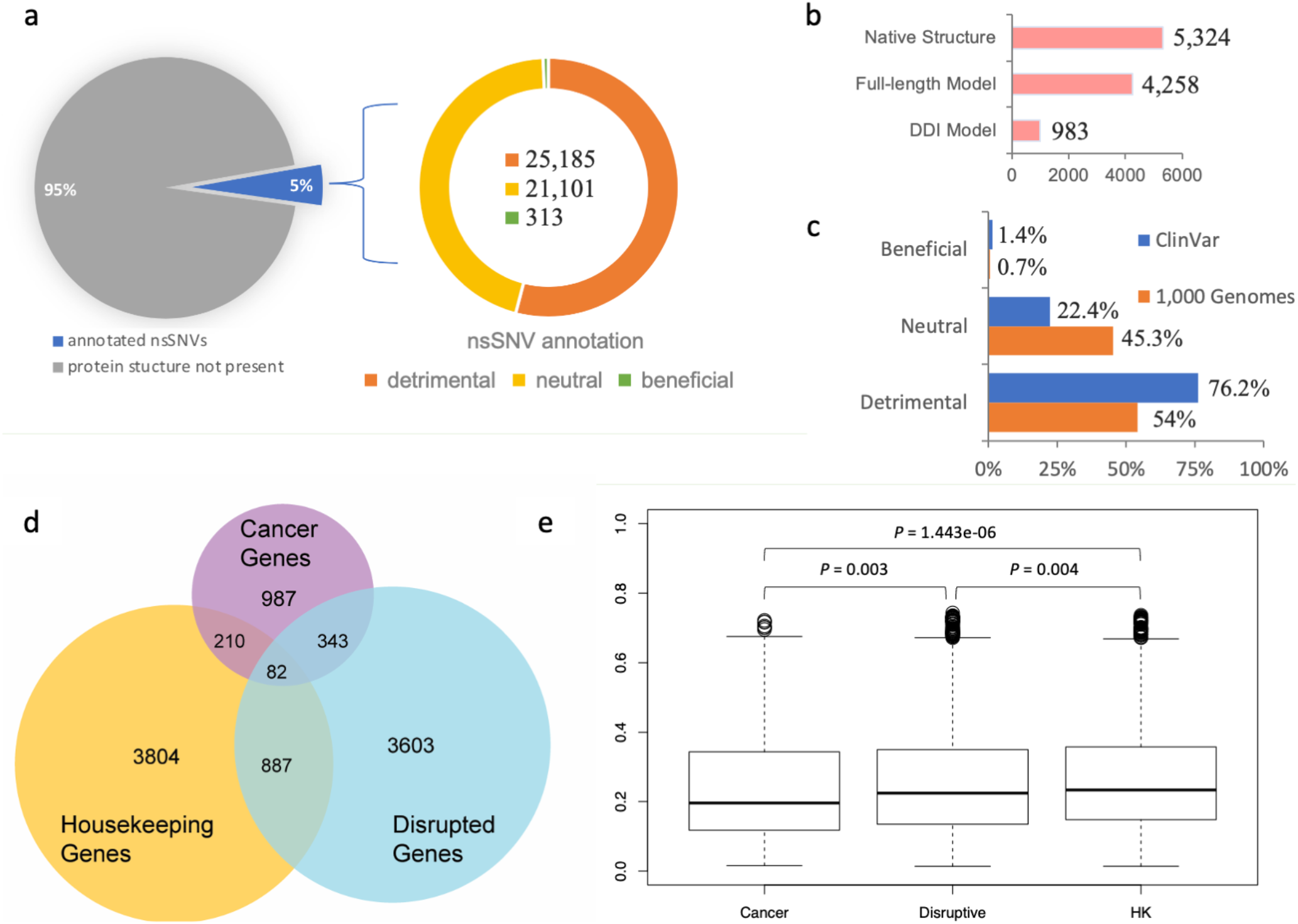
Analysis of population-specific nsSNVs annotated by SNP-IN tool. (a) SNP-IN tool annotation covers about 5% of total non-synonymous SNVs in the 1,000 Genomes Project; all three type of effects on PPIs were detected, with detrimental nsSNVs being the dominant type and beneficial nsSNVs contributing a substantially smaller fraction of variants than the other two types. (b) Three sources for the structural data on protein interaction complexes: native PPI structure, full-length PPI model, and domain-domain interaction (DDI) model. Overall, 46,599 nsSNVs mapped to 10,565 protein interaction complex structures were annotated; multiple nsSNVs were annotated by more than one source. (c) Comparison between SNP-IN tool based annotations of pathogenic mutations from ClinVar database and population-specific mutations from 1,000 Genomes Project. As expected, the population-specific nsSNVs obtained from the healthy population include a significantly larger proportion of neutral mutations compared to the pathogenic nsSNVs. (d) Three sets of genes used in our dN/dS analysis and their mutual overlap: disruptive genes, cancer genes, and housekeeping (HK) genes. (e) Comparison of *dN*/*dS* ratios across three gene sets.

Our results showed that, among the 46,599 population-specific nsSNVs (5% of all collected nsSNVs) (Fig. 2a), 25,185 nsSNVs (54.1% of annotated nsSNVs) were predicted as detrimental to at least one PPI involving the corresponding mutant protein, and only 313 nsSNVs (0.7% of annotated nsSNVs) were labeled as beneficial to a PPI (See Fig. 2c). In our previous work, we applied the same *in-silico* protocol to annotate a comprehensive set of 3,401 pathogenic mutations occurring in the genes associated with human diseases extracted from the ClinVar database [29]. We found that 2,592 nsSNVs (76.2%) were predicted as detrimental to at least one PPI, and 48 nsSNVs (1.4%) were predicted as beneficial (Fig. 2c). Our results showed that the proportion of the detrimental SNVs obtained from the 1,000 Genomes Project dataset, and thus observed in the healthy population, was 1.5 times smaller, but there was still a substantial number of nsSNVs that were expected to alter protein-protein interaction. Interestingly, both datasets of mutations shared significantly smaller fractions of the beneficial mutations, supporting our previous observation that genetic variations enhancing the protein-protein interaction were rare in the human genome [30]. An nsSNV predicted as detrimental is likely to be functional too, since it disrupts a PPI, and substantially affects the molecular function associated with it. If the loss of function further increases an individual’s susceptibility or predisposition to a certain disease, then the detrimental nsSNV is also a deleterious one. Interestingly, the abundance of the detrimental mutations found in our work was comparable with but slightly higher than the previously reported estimated overall abundance of the moderately to strongly deleterious mutations in a single genome, which ranged between 29% and 49% [31, 32].

We next studied the evolutionary conservation of genes harboring detrimental mutations. Specifically, we examined whether these genes tended to evolve at different rates compared to other well-characterized gene sets, such as cancer genes and housekeeping genes. To initiate this study, we defined “disruptive” genes as those carrying at least one deleterious nsSNV, as predicted by the SNP-IN tool. We then collected a cancer gene set from COSMIC [33] and MutPanning [34], resulting in 987 genes with high confidence. Next, we considered a set of 3,804 housekeeping genes (Fig. 2d, see *Methods*). After calculating the evolutionary rates (dN/dS) for each group (dN/dS median values for housekeeping, detrimental, and cancer genes sets were 0.235 ± 0.158, 0.223 ± 0.159, and 0.197 ± 0.157, respectively), we found that the differences between the dN/dS ratios of the three gene sets were all significant (Fig. 2e). Disruptive genes had a slighter higher dN/dS ratio compared to the cancer genes (P = 0.003). At the same time, disruptive genes were found to have a slightly lower ratio than the housekeeping genes (P = 0.004).

### Genes enriched with detrimental mutations are associated with diverse molecular functions and are implicated in various diseases

Following the observation of widespread disruptions in the human interactome caused by common variants occurring in the healthy populations, we focused on the genes whose protein products are enriched with detrimental mutations, which is a more constrained set compared to the set of disruptive genes introduced earlier. Genes enriched with detrimental mutations were defined as those with their disruptive mutation rate larger than the average rate plus the standard deviation (see Methods). Among all 3,603 proteins that carried nsSNVs with the functional outcome annotated by the SNP-IN tool, there were 461 proteins (12.8%) enriched with detrimental mutations.

To determine if the set of 461 genes enriched with detrimental mutations shared any functional patterns, we performed a GO enrichment analysis. We only selected the third-level GO terms in the GO hierarchy, which represented a reasonable trade-off between the level of detail each term could provide and how well that term was populated with the gene instances. In total, we extracted 458 GO terms with significant p-value after the multiple hypothesis correction (see Methods). Examining the significant GO terms revealed a few interesting findings. First, in the molecular function category, we obtained 62 GO terms enriched in the gene set with significantly high number of detrimental nsSNVs. The majority of the top 20 significant molecular function GO terms (See Supplementary Fig. S1) were related to molecular binding, including nucleotide binding (GO:0000166), TAP binding (GO:0046977), pyridoxal phosphate binding (GO:0030170). This implied that while the detrimental mutations were expected to exclusively disrupt the protein-protein interactions, the genes carrying them could be also involved in other molecular interaction mechanisms. In the biological process GO term category, more than one-third of terms were related to the immune system response. The immune system consists of many molecular interactions and processes implemented to protect the organism against a wide variety of pathogens and diseases. We note that, besides the occurrence of a significantly high number of detrimental mutations on these genes, there was also an uneven distribution of these mutations across different populations. The differences in the resulting network rewiring effect could be linked to the population phenotypic variance, such as population-specific disease susceptibility.

Generally speaking, such a disruptive effect on a protein-protein interaction is expected to alter or completely abolish the regular biological processes or molecular functions, potentially leading to a disease phenotype. Thus, we further investigated any potential links between these genes enriched with detrimental mutations and complex diseases. We examined the set of genes enriched with detrimental mutations using OMIM [35] and HGMD [36] databases. Indeed, while all these mutations originated from the genomes of healthy individuals, the 461 genes were associated with various complex diseases (See Supplementary Table S1).

### Edgotype analysis reveals distinct protein edgetic profiles in major populations

The human interactome, or the human protein-protein interaction network, is the set of protein-protein interactions that occur in a human cell. The interactome is represented as a graph, with nodes and edges corresponding to the individual proteins and their pairwise physical interactions, respectively. The edgetic network perturbation model, or “edgotype”, emphasizes the disruption of specific edges. However, unlike the traditional gene-centric model, which mainly focuses on the overactivity or silencing of gene expression, the edgotype can easily incorporate the influence of genetic variation.

Utilizing the edgotype concept, we systematically characterized the edgetic profile for the genes and their proteins enriched with detrimental mutations across different populations. To simplify the analysis, we treated the mutations with the neutral and beneficial effects as the same group of mutations that preserve the protein-protein interaction, as opposed to the interaction-disrupting effect of the detrimental mutations. We then excluded the interaction-preserving mutations from constructing the edgetic profile. Thus, the edgetic profile of a protein in a healthy population was represented as a sequence of binary vectors. Given a protein-protein interaction that the protein of interest was involved in and the protein’s binary vector, 1 corresponded to a detrimental mutation with non-zero allele frequency in the population, and 0 corresponded to a detrimental mutation that was not present in this population. The difference between the gene edgetic profiles in two different populations is quantified using the Manhattan distance between the extended vectors (see Methods). For each protein, we calculated the average pairwise edgetic profile difference between any two populations. This average difference was further normalized by the total number of detrimental mutations. Thus, the average difference value of 0.0 would correspond to a protein with an identical edgetic profile across all populations, indicating no differences in functional effects across the populations. On the other hand, the difference value of 1.0 would correspond to a protein with each detrimental mutation for each PPI present only in one population, indicating a unique combination of the interaction-disrupting mutations in each population. On average, the normalized difference of the total detrimental mutations occurring in one protein between any two major populations was 0.43. Such a distinct prevalence of detrimental mutations in different populations, combined with the unique interactions these mutations target, can help explain the genetic diversity and phenotypic variance across populations.

### Edgetic properties of population-specific nsSNVs could help explain the phenotypic variance across different populations

Our computational predictions enabled the characterization of the edgetic property of genetic mutations. More importantly, these functional characterizations of mutations, together with the population-specific genotype information, enabled us to generate a concrete “molecular mechanism” hypothesis for certain complex phenotypes and to explain the phenotypic variance across different populations. Our approach to analyzing the genotype-phenotype relationship could be well illustrated by an example of edgetic mutation rs671 on gene ALDH2. ALDH2 gene encodes aldehyde dehydrogenase 2, a member of a family of enzymes that metabolize alcohol and plays a major role in ethanol catabolism [37]. Another gene, ALDH1A1, encodes retinal dehydrogenase 1 which oxidizes retinaldehyde [38], but also has a broader specificity and oxidizes other aldehydes [38]. The missense rs671 mutation of ALDH2 gene was well known as the culprit in the phenomena of “Asian Flush” [39, 40], where an Asian person developed a flush reaction, turning red in the face, neck, and shoulders, after drinking alcohol [41, 42]. The frequency of individuals carrying rs671 was the highest in eastern Asia (MAF = 17.6%) but was almost absent among other major populations [41]. ALDH2 was involved in two protein-protein interactions: one self-interaction which was the basis for its homo-oligomerization, and another one with ALDH1A1. According to the SNP-IN tool prediction, rs671 could disrupt both of these interactions (Fig 3c). ALDH2 functioned as a homo-tetramer in the cell, and the rs671 allele resulted in a mutant ALDH2*2 protein which caused a critical disruption to ALDH2-ALDH2 interaction. Such disruption destabilized the tetramer, interfered with catalytic activity, and increased the protein turnover [37]. As a result, it made ALDH2*2 protein to be defective at metabolizing alcohol. It is worth noting that rs671 might also disrupt ALDH1A1-ALDH2 interaction, further reducing the ethanol catabolism. There was another mutation rs8187929 on gene ALDH1A1 that could exert a similar impact on ALDH1A1-ALDH2 interaction (Fig 3c). A recent study reported that rs8187929 was related to alcohol consumption and drinking behaviors in the Japanese population [43]. Similar to rs671, rs8187929 also predominantly existed in the East Asian population, but with a much lower allele frequency (MAF = 4.56% in East Asians). Importantly, while occurring in the same protein, the edgetic profile, which lists the interactions that were disrupted by the mutations, of rs671 was found to be clearly distinct from rs8187929. According to the edgetic profiling, rs8187929 had a more limited impact on the enzymatic activities involved in ethanol metabolism, potentially disrupting only one PPI, while rs671 had a substantially wider rewiring effect on the corresponding protein-protein interactions with a higher prevalence in the East Asian population.

**Figure 3.**
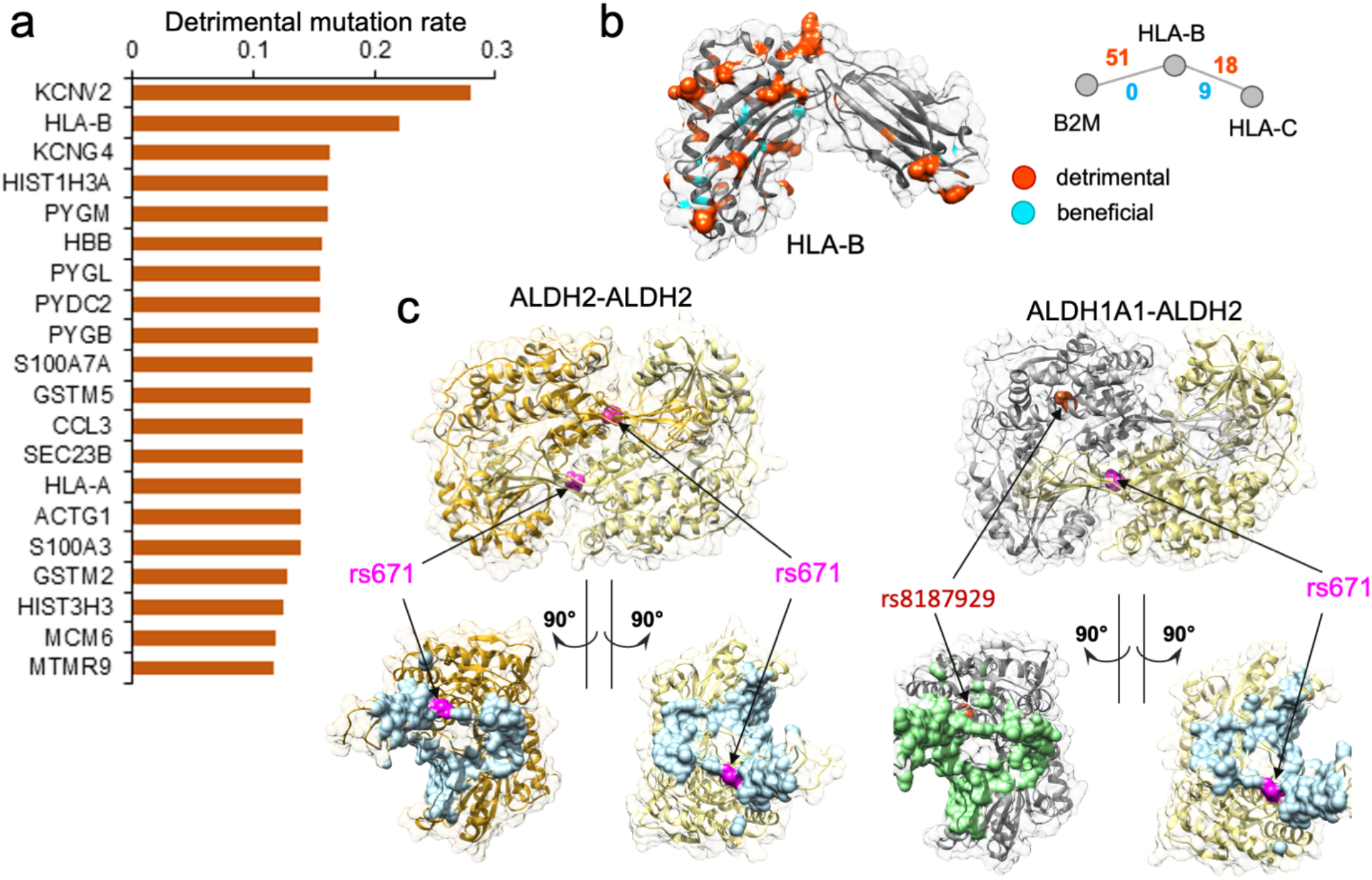
Genes enriched with detrimental mutations are associated with diverse molecular functions. (a) Top 20 genes with the highest disruptive mutation rate, normalized by protein sequence length. (b). Analysis of population-specific mutations affecting PPI mediated by HLA-B. HLA-B is one of the top genes with the highest rate of detrimental mutations regulating two PPIs, but also with a high number of beneficial nsSNVs. Shown is the protein structure of the HLA-B with the residues corresponding to detrimental mutations highlighted in orange and beneficial mutations highlighted in cyan. The two PPIs mediated by HLA-B and the number of detrimental and beneficial mutations associated with each PPI. Detrimental mutations are widespread on the structure. (c) Case study of two protein-protein interactions implicated in the “Asian Flush” phenotype. Shown are two mutations: rs671 and rs8187929 mapped w.r.t protein binding interface colored in light blue for ALDH2 and light green for ALDH1A1. rs671 is predicted by SNP-IN tool to disrupt the two PPIs. While rs8187929 is also related to human drinking behavior, it may be less effective than rs671.

We performed another in-depth study on HLA-B gene, which was among the genes with the highest disruptive mutation rate, 22.1% (Figs 3a, 3b). HLA-B (major histocompatibility complex, class I, B) is a human gene encoding a protein that plays an important role in the immune system, helping distinguish the organism’s native proteins from the foreign ones [44], and enabling the immune system to react to a broad spectrum of pathogens [45–48]. Together with two other related proteins, HLA-A and HLA-C, HLA-B belongs to the major histocompatibility complex (MHC) gene family. Interestingly, HLA-A was also found among the top 461 genes enriched with detrimental mutations (Supplementary Fig. S2). The high number of normal variations disrupting protein-protein interactions suggested that diverse immune responses to various pathogens could be partially attributed to many distinct HLA gene edgetic profiles and the unique rewiring subnetwork centered around the HLA family. We hypothesize that, in conjunction with the unequal distribution of mutation frequency across populations, the rewiring effect of HLA complex could contribute to the different disease susceptibility and the varied immune response.

### Comparative network analysis shows disease mutations target less efficient subnetworks

We next compared the topological properties and the rewiring effects in the human interactome of both the pathogenic mutations curated from the ClinVar database and the population mutations from this study. First, we constructed the human interactome by utilizing two main protein-protein interaction data sources, HINT (High-quality INTeractomes) and HuRI (Human Reference Interactome). The unified human interactome consisted of 105,087 interactions (see Methods). From the list of human disease genes retrieved from the ClinVar database, 576 genes carried at least one detrimental mutation, and the detrimental mutations occurring on these pathogenic genes rewire 1,162 interactions. There were 3,603 genes with at least one detrimental mutation from the 1,000 Genomes Project with the number of rewired interactions equal to 4,529. The overlaps between the two gene sets and the two interaction sets were 449 genes and 736 interactions, respectively. The overlaps of genes and the affected interactions were relatively high, suggesting that both pathogenic mutations and normal variants could cause detrimental effects on the same subset of interactions (Fig. 4a). We, thus, hypothesized that the presence of detrimental mutations in healthy populations might not be sufficient to cause the disease phenotype, but could nevertheless contribute to the disease susceptibility.

**Figure 4.**
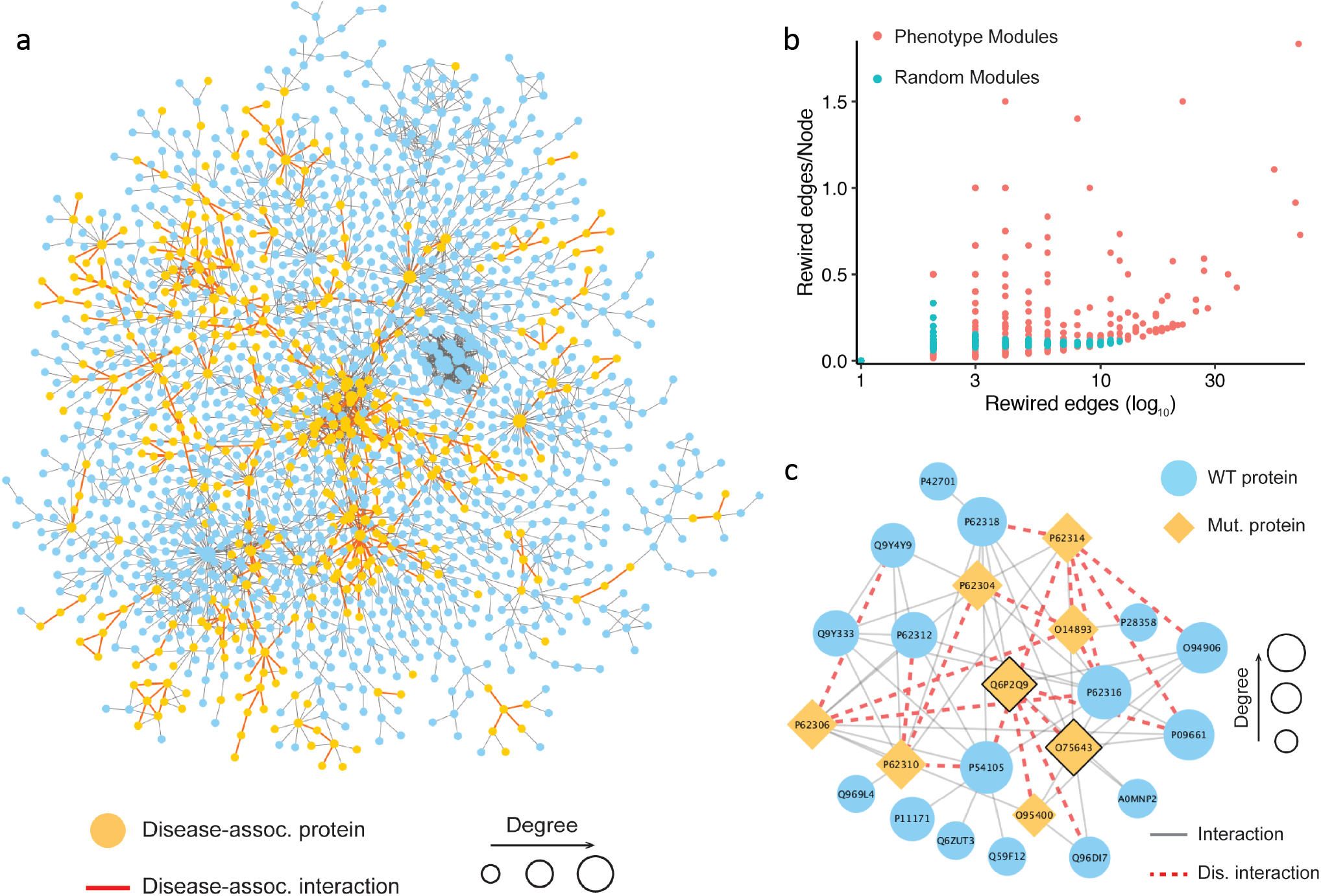
Analysis of phenotype-associated modules detected based on the topological properties. (a) Network visualization of the largest connected component targeted by the pathogenic mutations and common mutations. The yellow node represents a gene carrying pathogenic mutations, and the red edge represents an interaction targeted by both the pathogenic mutations and common mutations. (b) Comparison of interaction disruptions accumulated in the phenotype associated modules and random modules. The results show that a phenotype-associated module carries more detrimental mutations than a random one. (c) An example of the phenotype-associated module in the human interactome detected based on the DSD idea. Shown is a dense PPI module with the mutant proteins carrying variations disrupting PPI interactions, indicated by the orange nodes for the proteins and red dashed edges for the PPIs, respectively.

We next considered two groups of disrupted interactions caused by nsSNVs in the context of the whole human interactome and examined their topological properties. To account for the exclusive effects elicited by each set of the mutations, we excluded the 736 interactions shared between the two datasets. Then, we measured the centrality of the interactions rewired by the two groups of mutations using two main network edge centrality measures, the shortest-path edge betweenness and the flow edge betweenness (see Methods). Our results showed that there was no significant difference between the two disrupted interaction sets in terms of both edge centrality measures. Furthermore, the results showed that both, the pathogenic mutations and the mutations found in healthy populations, did not tend to disrupt interactions of high centrality, thus avoiding critical breakdowns in the interactome. We further investigated the network rewiring effect from a more indirect perspective, by collapsing the two disrupted interaction sets into unified subnetworks and examining their network efficiency, a measure that described how efficiently a network exchanged the information. Because protein-protein interaction networks have been previously determined to be a small-world network [49], the network efficiency is generally better preserved during the randomized attack, compared to the one targeting the hub proteins [50]. We found that the subnetwork disrupted by the pathogenic mutations had a lower network efficiency compared to the population-specific mutations, with a nearly two-fold difference (0.0117 vs. 0.0223). The fact that the pathogenic mutations target less efficient networks than the population mutations, suggests that they occur, en masse, in more random positions of the interactome, while the occurrence of the population mutations appears to be more controlled and evolutionary guided.

### Phenotype-associated modules in the human interactome are enriched with detrimental mutations

Modular structure is one of the essential characteristics of biological networks [51]. The identification of these modules is a crucial step in the network analysis helping to uncover the biological mechanisms underlying complex phenotypes [20, 52]. To identify biologically relevant modules in the human interactome, we applied two distinct module detection methods: the first method relied only on the network’s topological information, with no prior biological knowledge, while the second method was a seed-based approach that leveraged the knowledge about the disease-associated proteins (see Methods). Given its outstanding performance in DREAM challenge [53], we expect for the first method to accurately discover the modules that are significantly associated with complex traits, including both healthyl phenotypes and disease phenotypes. For the second, biological knowledge-driven, approach, we adopted DIAMOnD [54]. The DIAMOnD method aimed to identify the full disease module around a set of known disease proteins (see Methods).

Using the first approach, we collected 1,055 possible phenotype-associated modules with high confidence with the module sizes ranging between three and 100 genes. We further investigated whether the PPI disruptions and rewiring inside these phenotypes associated modules caused by detrimental mutations had any role in developing a complex trait. To do so, we randomly generated the same number of modules and compared the number of rewired edges between the two module sets. Then, the number of the rewired edges in the module was normalized by the total number of nodes in the module. The results showed that the phenotype-associated modules had significantly more detrimental mutations than the random ones (Fig 4b). This finding supported our hypothesis that the rewiring caused by the detrimental mutations inside the phenotype-associated modules may directly contribute to the development of certain phenotypes. Shown in Fig. 4c is an example of detected module that includes 24 genes. The literature search confirmed that this module was associated with pre-mRNA splicing. Specifically, some genes in the module, such as PRPF8 (Uniprot: Q6P2Q9), GEMIN2 (Uniprot: O14893), and SNRNP200 (Uniprot: O75643), were the key components of the spliceosome, a complex molecular machine consisting of small nuclear RNAs (snRNAs) and approximately 80 proteins [55]. In this module, the detrimental mutations targeted interactions mediated by two protein hubs, PRPF8 and SNRNP200 (Fig. 4). PRPF8 forms a scaffold to help in the assembly of the snRNAs and proteins of the complex, and SNRNP200 mediates the interaction of snRNAs and catalytic activity. Mutations in these proteins have been previously linked to a number of traits [56–59]. These traits include both normal phenotypes and disease phenotypes, such as leukocyte count, body mass index, balding measurement, coronary artery disease, and others. Although the phenotypic nature of the interaction-disrupting variants has not been determined, our results provide mechanistic insights into functional variation by pinpointing specific protein-protein interactions.

We next carried out seed-based module detection on the human interactome. To do so, we first collected disease-associated genes as seeds for 70 complex diseases. We note again that these mutations originated from the healthy populations hence the interaction disruptions caused by them were not expected to cause severe functional changes in the cell. However, we suspected that certain population-specific changes in these disease-associated modules caused by detrimental mutations could be linked to the differences in disease susceptibility across different ethnic groups. We then run the DIAMOnD algorithm with the disease-associated genes as partial input. For the 70 disease modules collected from DIAMOnD output, we summarized the network-aggregated results in the form of a heatmap (Fig 5a). In the heatmap, the disease-associated modules and the populations were represented by rows and columns, respectively. The color of each cell in the heatmap represented the prevalence of rewiring in a specific disease module and a population, where red represented a higher occurrence of disrupted interactions in the module. To uncover patterns, we also applied hierarchical clustering to group diseases and populations by the similarity of rewiring prevalence. As a result, we found that among the five major populations, detrimental mutations and the resulting PPI rewiring were most prevalent in East Asians and Africans across 70 disease modules.

**Figure 5.**
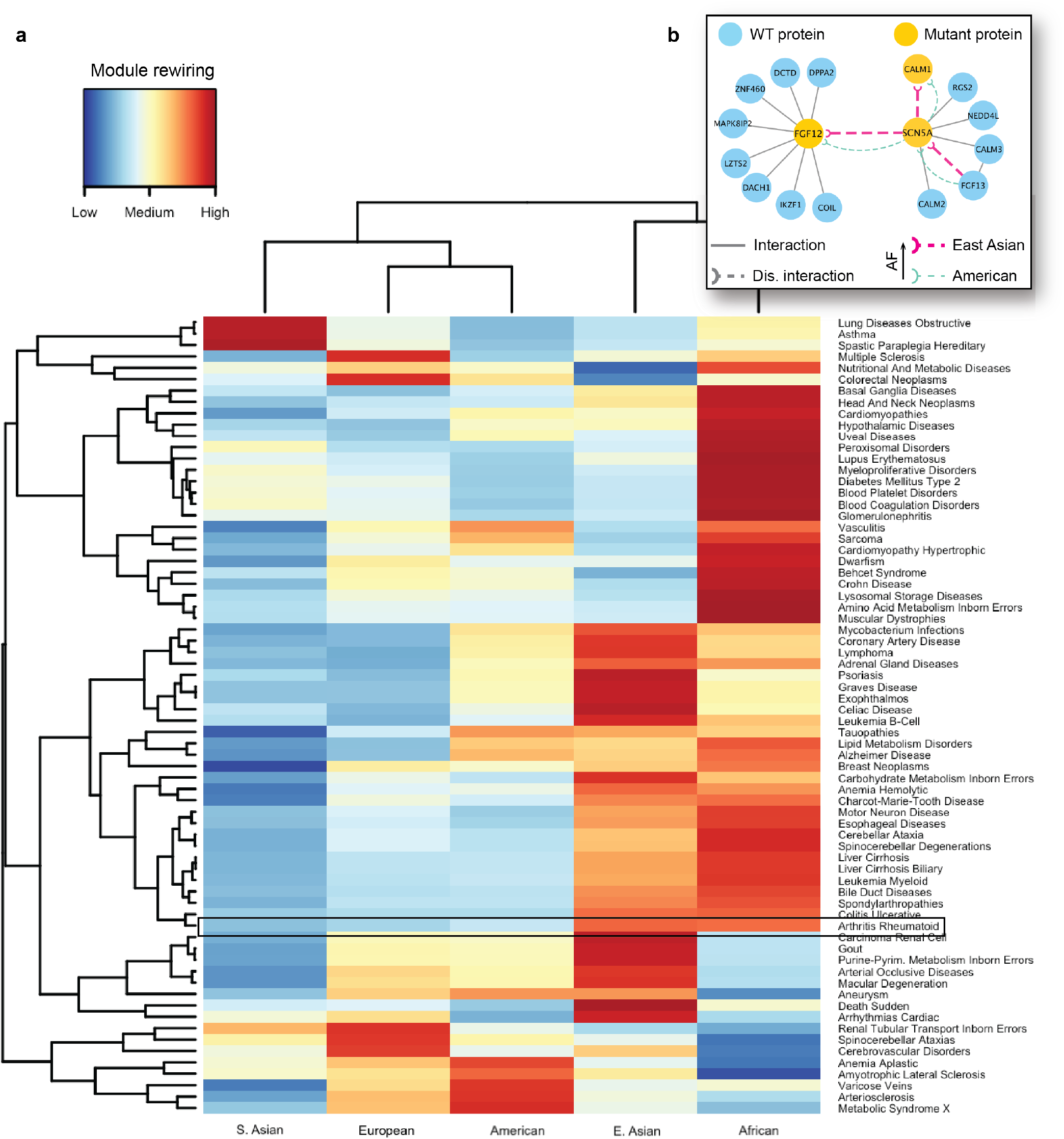
Analysis of disease modules impacted by detrimental mutations. (a) A heatmap showing the different prevalence and disruption levels caused by the population-specific mutations across different populations. The disease-associated modules are represented by rows and the populations are represented by columns. The color of each cell in the heatmap represents the prevalence of rewiring in a specific disease module for a given population. (b) Key interactions damaged in the module associated with arrhythmias carry the detrimental mutations with different prevalence in East Asian and Americans. FGF12 and SCN5A are two important genes implicated in arrhythmias, and disruptions associated with FGF12 and SCN5A are more frequent among East Asians compared to the American population.

As an in-depth study, we investigated the disease module for cardiac arrhythmia (Supplementary Fig. S3). A cardiac arrhythmia is a group of conditions that cause the heart to beat too fast, too slow, or irregular in timing [60]. Arrhythmia affects millions of people and usually develops in the older adults, with many affected people not requiring medical attention or treatment. However, this condition can also have lethal outcomes, such as sudden cardiac death. Even though there are far fewer cardiac genetic studies in Asian populations than in Western populations, a recent review on arrhythmias [61] found that Japanese individuals carry a higher prevalence of long QT syndrome and Brugada syndrome, which can both increase the risk for sudden cardiac death. Several well characterized proteins associated with arrhythmias are known to interact with each other. A protein-protein interaction associated with Brugada syndrome and sudden death is formed between FGF12 (Fibroblast growth factor 12) and SCN5A (sodium voltage-gated channel alpha subunit 5) proteins. Fibroblast growth factor homologs (FGF11–FGF14) perform many intracellular functions and are well known to bind to sodium and calcium channels, modulating cardiac currents. Among them, FGF12 is the major fibroblast growth factor expressed in the human cardiac ventricle. Another key protein, SCN5A is a sodium channels that plays an important role in modulating electrical impulses and their conduction in the heart. SCN5A mutations [62] are implicated in many arrhythmias such as long QT syndrome, Brugada syndrome, atrial fibrillation, progressive cardiac conduction defect, and sick sinus syndrome. In addition, SCN5A is known to interact with the calmodulin gene products (CALM1, CALM2, and CALM3) that are also associated with susceptibility to arrhythmias [63].

As indicated by the heatmap, there was a substantial difference in the overall PPI rewiring prevalence between the East Asian and American populations (Fig. 5a). When we selected the disrupted interactions of each of these populations and mapped them to the arrhythmia-associated module (Fig. 5b), we saw distinct rewiring patterns in the PPIs associated with arrhythmias in East Asians and Americans. We further focused on a few specific interactions mediated by the aforementioned arrhythmia-associated proteins (Supplementary Fig. S3). For the interaction between FGF12 and SCN5A, we predicted the rewiring due to mutations in FGF12, in which the deleterious variant is more frequent in East Asians than in Americans (Fig. 5b, thick and thin edges respectively). This PPI disruption by the mutant proteins had been experimentally studied in mice and humans [64]; it showed that such disruption can cause a sodium channel loss-of-function phenotype. More specifically, experiments revealed the reduced binding of the mutant FGF12 to SCN5A and the resulting reduction in sodium channel current which affected the cardiac ventricular action potential. Next, our analysis showed that the disruption (Supplementary Fig. S3) between SCN5A and CALM1 was also more frequent in the East Asian population. Finally, we predicted another interaction of SCN5A, with FGF13, to be disrupted, due to a detrimental variant in the former protein. The SCN5A-FGF13 PPI was also implicated in arrhythmias [65]. This analysis provided us with the mechanistic explanation of the molecular mechanisms driving the increased susceptibility of the disease in the selected populations.

## DISCUSSION

In this work, we created a comprehensive catalog of population-specific edgetic effects at the whole-interactome level. Our work leveraged a recently developed novel machine learning approach, the SNP-IN tool, that determined the interaction-rewiring effects of nsSNVs. This method was previously applied to variants associated with diseases, showing the feasibility of a large-scale *in-silico* edgetic study. To compare the functional impact of the population-specific variations, we applied our approach to annotate the protein-rewiring effects of a comprehensive set of 46,599 nsSNVs collected from the 1,000 Genomes Project. The functional impact of a variant on a PPI was annotated as neutral, detrimental, or beneficial based on their effect on the PPI quantified by a change in the binding affinity. In summary, we determined that 25,185 SNVs (54%) were predicted as detrimental to at least one PPI, indicating that a significant number of nsSNVs from the healthy populations can alter the protein function. Our findings on the dN/dS ratios of the disruptive gene set showed a slight difference of the ratios between the disruptive gene set and two other sets, the housekeeping genes and cancer drivers, both of which are known to be evolutionary conservative. The differences are statistically significant, suggesting that the disruption of protein-protein interaction caused by nsSNVs imposes unique functional constraints on the human genome evolution. We also note that the differences are small, so one should use caution when making conclusions from the analysis.

While the traditional “node-centric” gene removal approach has been widely used to quantify the disruption in the network due to the gene loss, the edgotype approach provides a much finer-grain view of the mutation’s functional impact by focusing on alterations in specific (not necessarily all) molecular interactions mediated by the mutated protein. Indeed, many mutations have been shown to be edgetic [24, 30]. Besides, the interactions mediated by the same protein may not necessarily occur at the same time. Thus, the mutations accumulating on the same protein might affect different PPIs at different times. Edgetic network perturbation models have also been proposed and adopted to study complex genetic diseases [24, 25, 27, 30, 66]. Sahni *et. al.* suggested a relationship between the edgetic perturbations and disease severity [24]. Importantly, the edgotype provided a plausible explanation for some complicated genetic phenomena, such as locus heterogeneity and gene pleiotropy [27, 67]. In addition, the edgetic perturbation model offered a network-based hypothesis to explain modes of inheritance [25]. While all these studies have been focused on the disease-associated mutations, the current work suggests that the edgotype concept can be extended to population genetics to study the mutation frequency patterns and their functional impacts across healthy populations. Case studies in this work provide evidence that this edgotype-based approach can better explain the phenotypic variance across populations.

Our analysis revealed that genes enriched with detrimental mutations were linked to various disease phenotypes. Because these mutations were harbored among the healthy individuals, the mutations’ interaction-disrupting effects alone would not be enough to result in a disease phenotype. Instead, a distinct group of mutations, absent in the healthy population but present in the same genes, might inflict more severe damage to the molecular mechanisms connected with the disease. Nonetheless, the detrimental mutations in the healthy population could add to the cumulative damage of a protein-protein interaction, thereby potentially increasing susceptibility to the disease. This hypothesis received further supporting evidence from our comparative network analysis concerning disease mutations and normal mutations. This analysis shows that some of the healthy population mutations can target the same set of interactions that were previously shown to be targeted by the pathogenic mutations, leading to similar alterations in the network topology.

Finally, our research reveals a profound implication of detrimental mutations and their effects on network rewiring. We observed distinct population-specific rewiring patterns within disease modules, pointing to inherent differences in disease susceptibility across populations. Delving deeper into the network disease modules, our analysis uncovered that modules associated with specific phenotypes had a higher concentration of detrimental mutations compared to random modules. Moreover, when examining the effects of these mutations across different populations, the rewiring patterns within the disease-related network modules frequently emerged as unique to one or several particular populations. Overall, our large-scale *in-silico* approach sheds light on the mechanistic nuances of population-specific differences, enhancing our understanding of both health and disease traits.

## METHODS

### Genetic mutation data processing and construction of the human interactome

The mutation data are collected from the 1,000 Genome Project [5], a major international consortium effort to catalog genetic variants across major ethnic populations. In the functional annotation and downstream analysis, we focus exclusively on non-synonymous missense single nucleotide variants (nsSNVs) and excluded non-coding variations, synonymous SNVs, short indels, and structural variations. A single nucleotide variant is the simplest and yet the most common type of genetic variation among people. In this work, nsSNVs are selected because they are most likely to make a functional impact on the PPIs. The mutation data are first processed with ANNOVAR [68] to obtain SNV locations on the genes and their protein products and determine the corresponding residue substitution. This information is essential for the SNP-IN tool [16] to predict the rewiring effects of mutations.

To construct a unified human interactome, we use two large-scale protein-protein interaction data sources: High-quality INTeractomes database (HINT) [69], and Human Reference Protein Interactome Mapping Project (HuRI) [70]. HINT is a centralized database of human PPIs integrated from several other sources and annotated using both an automated protocol and manual curation. HuRI focuses on the experimentally validated PPIs using yeast-two-hybrid experiments. The two PPI sources are merged because they provide complementary views of the whole human interactome. The HINT database contains 63,684 interactions, while the HuRI dataset includes 76,537 interactions. When merged, the final human interactome includes 105,087 interactions, with 35,134 interactions occurring in both datasets.

### Functional annotation of nsSNV

Next, we determine if a mutation occurring in a protein can affect a protein-protein interaction that this protein is involved in. To do so, consider two PPIs: a wild-type interaction, which involves a wild-type version of that protein, and a mutant PPI, which involves the protein with the mutation. To accurately determine the functional damage to a PPI caused by a mutation, our approach requires information about both the mutation and the PPI structure or structural model as the input for the SNP-IN tool [16, 30]. The prediction task by the SNP-IN tool is then formulated as a classification problem, with three classes of rewiring effects: beneficial, neutral, and detrimental. The effects are assigned based on the difference between the binding free energies, ΔΔG, of the mutant and wild-type PPIs. Specifically,

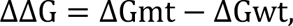

where ΔGmt and ΔGwt are the mutant and wild-type binding-free energies, respectively. The beneficial, neutral, or detrimental types of mutations are then determined by applying two previously established thresholds to the ΔΔG values [71, 72]:

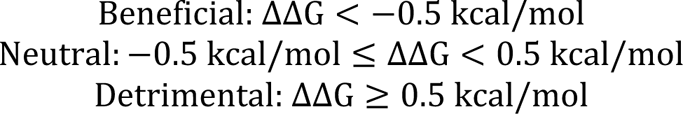

Obtaining a PPI structure for the SNP-IN tool is done in two ways. First, if a PPI has a native structure of the corresponding protein complex, we extract it from the Protein Data Bank [73]. If there is no native structure for a PPI, we apply homology, or comparative, modeling [74] to get a structural model, either for the full-length PPI complex or at least for a pair of protein domains that form the interaction interface [16, 30]. Lastly, we define a disruptive gene as a gene carrying at least one detrimental mutation, which causes a disruptive effect on the related protein-protein interactions based on SNP-IN tool output.

### dN/dS ratio calculation and comparison

The high-confidence collection of cancer genes is obtained by merging the Cancer Gene Census dataset [75] and a recently published computationally predicted cancer gene set, MutPanning [34]. The Catalogue of Somatic Mutations in Cancer (COSMIC) and Cancer Gene Census (CGC) are the ongoing efforts to catalog genes with known mutations that have been causally implicated in cancer. CGC is used as a golden standard in cancer genetics [75], with each driver gene being accompanied by an expert-curated description. MutPanning is a computational method for the identification of cancer driver genes across 28 tumor types that combines the characteristic contexts around passenger mutations with the signals of mutational recurrence [34]. The two cancer driver gene sources contain 723 and 460 genes respectively, with an overlap of 196 genes. In total, 987 distinct cancer genes are collected. Next, the list of 3,804 housekeeping genes is retrieved from a dataset compiled by Eisenberg and Lavanon [76]. Housekeeping genes are the genes that are essential for the existence of a cell and the maintenance of the basal cellular functions [77]. The genes are believed to express in all cells of an organism under normal and pathophysiological conditions [78].

The evolutionary analysis of the genes is carried out by calculating a ratio between the number of non-synonymous substitutions (dN) to the number of synonymous substitutions, dN/dS [79]. We calculate the dN/dS ratio by comparing the human gene sequences with the orthologous groups. Specifically, the orthologous genes of macaque, gorilla, orangutan, chimpanzee, and gibbon corresponding to the human genes under consideration (GRCh38.p13) are queried from the Ensembl Biomart platform [80] together with the corresponding dN and dS values. The dN/dS ratio is then calculated, while the missing and infinite values are removed. For sets of homologs that do not include exactly one representative in each organism, the group mean of ER is calculated as the representative. The overall ER of a human gene is the average value of ERs for an individual organism, and we only include genes with orthologs in more than three organisms. The comparison of ER between different sets of genes is done with Wilcoxon test because no prior information about the underlying distribution is known.

### Calculation of disruptive mutation rate in proteome and GO enrichment analysis

The disruptive mutation rate of a gene is the ratio of the total number of detrimental mutations in a corresponding protein to the protein’s length. We first collect all detrimental mutations occurring in the protein based on our SNP-IN tool annotation. The protein sequence length information is retrieved from Uniprot [81]. The average disruptive mutation rate across the proteome is considered as the background rate, and we define proteins (and the corresponding genes) with the disruptive mutation rate greater than the average rate plus a standard deviation to be enriched with the detrimental mutations.

We then investigate the biological implications of genes enriched with detrimental mutations using Gene Ontology (GO) [82]. For the set of genes enriched with detrimental mutations, a GO enrichment analysis is carried out to obtain a list of enriched GO terms. In the analysis, we use the third level of the GO hierarchy and consider only those GO terms with P-value ≤ 0.01. By comparing the results from the second and fourth levels, we find that the third level represents the best trade-off between having too general (and thus not very informative) but well-populated GO terms from the second level, and more specific but not well-populated (and thus not suitable for the enrichment analysis) terms from the fourth level. The GO enrichment is performed using DAVID [83], and multiple testing correction is done with the false discovery rate estimation [84].

### Population-specific edgetic profiles of genes enriched with detrimental mutations

For a protein (and the corresponding gene) enriched with detrimental mutations, we introduce the population edgetic profile concept to describe the diverse network rewiring effects caused by detrimental mutations centering around the mutated gene across different populations. Simply put, the edgetic profile, *EP*(*p*), of a protein *p* that participates in *m* interactions is represented as a sequence of vectors (*v_1_, v_2,_ …, v_m_*) of different lengths, *l_1_,l_2,_ …, l_m_*. Each vector *v*_*i*_ consists of a list of *l*_*i*_ detrimental mutations targeting the same interaction *i*. The element takes only binary values, where 1 indicates a detrimental mutation with a non-zero allele frequency for a specific population, and 0 indicates a detrimental mutation that is not present in this population. For example, *EP*(*p*) = ([1, 0, 0, 1], [0, 1, 1], [1, 0, 0, 1, 1]) corresponds to an edgetic profile of a protein that has three interaction partners.

To compare the edgetic profiles, *EP*^(*i*)^ and *EP*^(j)^, of the same protein in two different populations, *i* and *j*, respectively, we first calculate the Manhattan distance between two profile vectors after merging the original sequence of vectors into a single vector of dimension 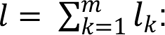

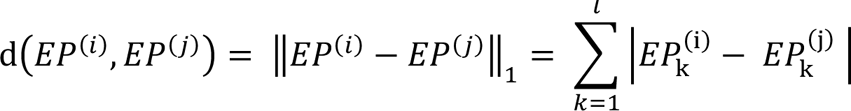

The Manhattan distance between two edgetic profiles is further normalized by taking the total number of detrimental mutations into account:

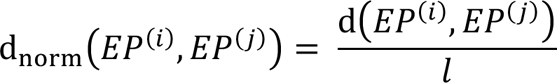

The average difference of gene edgetic profiles between any two populations is then evaluated as follows:

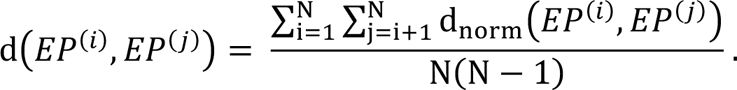

### Examination of topological properties of genes enriched in detrimental mutations in human interactome

In our comparative network analysis, we independently investigate and compare the topological importance of the rewired edges targeted first by the pathogenic mutations and then by the healthy population mutations. To quantify the importance, we apply two edge centrality measures to the network: shortest-path edge betweenness and current-flow betweenness. Betweenness centrality was proposed as a general measure of centrality [85]. It was applied to a wide range of real-world problems, including problems related to biological [86], social [87], and transportation networks [88]. Typically, an edge with a higher betweenness centrality in a complex network would be more critical for the network, because more information would pass through that edge. In the human interactome, mutations disrupting an interaction corresponding to such an edge are likely to have a greater impact on the cell functioning. The shortest path edge betweenness centrality is a measure of centrality based on the number of the shortest paths that go through a given edge. The measure is defined as the sum of the fractions of all-pair shortest paths passing through an edge [89]. Formally, the shortest path edge betweenness centrality of an edge *e* is given by the expression:

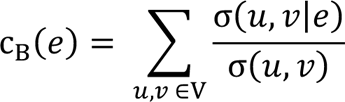

where V is the set of nodes, σ(*u*, *v*) is the number of shortest paths between a pair of nodes, *u* and *v*, and σ(*u*, *v*|*e*) is the number of those paths passing through edge *e*. The current-flow betweenness is another global centrality measure, which is based on an electrical current model for information dispersion. It is also known as random-walk betweenness centrality [90].

Another graph-based metric to characterize the rewiring effect caused by detrimental mutations is the network efficiency [91]. The concept of efficiency measures how well the network propagates and exchanges information. To compute the efficiency of the subnetworks targeted by pathogenic mutations and normal mutations, we first constructed two separate subnetworks by grouping the corresponding disrupted interactions. For a pair of nodes in the network, the efficiency is the multiplicative inverse of the shortest path distance between the pair. The global efficiency of a graph is defined as the average efficiency of all pairs of nodes. Formally, it is defined as:

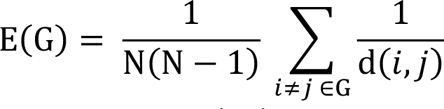

where N is the number of nodes in a network and d(*i*, *j*) is the length of the shortest path between a pair of nodes, *i* and *j*.

### Phenotype-associated community detection in human interactome and its enrichment with detrimental mutations

Discovering biologically relevant modules is a challenging task [92, 93]. Methods for module discovery primarily come in two types. Methods of the first type identify the modules in a biological network using exclusively the information on the network’s topology. The main challenge for such methods is the lack of any relevant biological information about the proteins constituting the network. Methods of the second type initiate the module with the “seed” genes or proteins, and then gradually grow the module by attaching additional genes/proteins from the network. In this work, we explore methods of both types.

For the first type, we adopt a technique based on the idea of “diffusion state distance” (DSD) [94], which has been proven to be the best performer in the DREAM Module Identification challenge [53], as it yields the highest number of the phenotype-associated modules. The approach has two main steps: computing a DSD matrix and applying spectral clustering on the DSD matrix to identify phenotype-associated modules. As a proximity measure, DSD is conceptually different from the traditional shortest-path measure: the latter favors the hub-like nodes in the network and does not incorporate information about the intrinsic network structure. Intuitively, a protein pair connected by paths through the low-degree nodes share more functional similarity than other protein pairs connected by paths that go through the hubs. (See Supplementary Fig. S4). So, DSD is a more fine-grained measure of similarity that “downweighs” the hubs in the human interactome. Formally, given an undirected graph *G*(*V*, *E*) consisting of a node set *V* = {*v*_1_, *v*_2_, …, *v*_n_} and |*V*| = *n*, we define a vector *D*^k^(*A*):

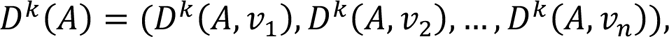

where *D*^k^(*A*, *v*_i_) is the expected number of times that a simple symmetric random walk starting at node

*A*, and proceeding for *k* steps, will visit node *v*_$_. Thus, *D*^k^(*A*) defines a global distance measure from node *A* to all the other nodes of the network. Assume *k* is fixed. Then, the DSD between nodes *A* and *B* is defined as follows:

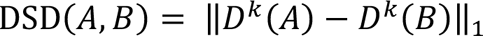

The DSD matrix is calculated using the “cDSD’’ method [95] from software available at: http://dsd.cs.tufts.edu/capdsd. The follow-up spectral clustering on the DSD matrix is performed with scikit-learn package [96].

For the second type, we consider a recently published method, DIseAse MOdule Detection (DIAMOnD) [54]. DIAMOnD is a module detection algorithm that utilizes known seed genes to identify disease modules by adding new proteins according to their connection significance to the seed proteins. The algorithm exploits the fact that the disease-associated proteins do not reside within locally dense communities, and thus the significance of their connections would be a more predictive measure when compared to the local network density. In addition, the use of the significance of the number of connections reduces spurious detection of the high-degree proteins, compared to using the absolute number of connections. As a result, DIAMOnD produces a connected disease module together with a list of disease-associated protein candidates ranked by the connectivity significance.

## REFERENCES

1. Schuster, S.C., Next-generation sequencing transforms today’s biology. Nature methods, 2008. 5(1): p. 16–18.

2. Metzker, M.L., Sequencing technologies—the next generation. Nature reviews genetics, 2010. 11(1): p. 31–46.

3. Shendure, J., et al., DNA sequencing at 40: past, present and future. Nature, 2017. 550(7676): p. 345–353.

4. Consortium, I.H., The international HapMap project. Nature, 2003. 426(6968): p. 789.

5. Consortium, G.P., A global reference for human genetic variation. Nature, 2015. 526(7571): p. 68–74.

6. van Rooij, J.G., et al., Population-specific genetic variation in large sequencing data sets: why more data is still better. European Journal of Human Genetics, 2017. 25(10): p. 1173–1175.

7. Consortium, G.P., An integrated map of genetic variation from 1,092 human genomes. Nature, 2012. 491(7422): p. 56.

8. Sherry, S.T., et al., dbSNP: the NCBI database of genetic variation. Nucleic acids research, 2001. 29(1): p. 308–311.

9. Alexander, R.P., et al., Annotating non-coding regions of the genome. Nature Reviews Genetics, 2010. 11(8): p. 559.

10. Cui, H., et al., The variation game: Cracking complex genetic disorders with NGS and omics data. Methods, 2015. 79: p. 18–31.

11. Pabinger, S., et al., A survey of tools for variant analysis of next-generation genome sequencing data. Briefings in bioinformatics, 2014. 15(2): p. 256–278.

12. Cooper, G.M. and J. Shendure, Needles in stacks of needles: finding disease-causal variants in a wealth of genomic data. Nature Reviews Genetics, 2011. 12(9): p. 628–640.

13. Ward, L.D. and M. Kellis, Interpreting noncoding genetic variation in complex traits and human disease. Nature biotechnology, 2012. 30(11): p. 1095–1106.

14. Raphael, B.J., et al., Identifying driver mutations in sequenced cancer genomes: computational approaches to enable precision medicine. Genome medicine, 2014. 6(1): p. 5.

15. Cline, M.S. and R. Karchin, Using bioinformatics to predict the functional impact of SNVs. Bioinformatics, 2011. 27(4): p. 441–448.

16. Zhao, N., et al., Determining effects of non-synonymous SNPs on protein-protein interactions using supervised and semi-supervised learning. PLoS computational biology, 2014. 10(5): p. e1003592.

17. Barabasi, A.-L. and Z.N. Oltvai, Network biology: understanding the cell’s functional organization. Nature reviews genetics, 2004. 5(2): p. 101–113.

18. Zhang, B., Y. Tian, and Z. Zhang, Network biology in medicine and beyond. Circulation: Cardiovascular Genetics, 2014. 7(4): p. 536–547.

19. Carter, H., M. Hofree, and T. Ideker, Genotype to phenotype via network analysis. Current opinion in genetics & development, 2013. 23(6): p. 611–621.

20. Ideker, T. and R. Sharan, Protein networks in disease. Genome research, 2008. 18(4): p. 644–652.

21. Sahni, N., et al., Edgotype: a fundamental link between genotype and phenotype. Current opinion in genetics & development, 2013. 23(6): p. 649–657.

22. Zhong, Q., et al., Edgetic perturbation models of human inherited disorders. Molecular systems biology, 2009. 5(1): p. 321.

23. Dreze, M., et al., ’Edgetic’perturbation of a C. elegans BCL2 ortholog. Nature methods, 2009. 6(11): p. 843.

24. Sahni, N., et al., Widespread macromolecular interaction perturbations in human genetic disorders. Cell, 2015. 161(3): p. 647–660.

25. Zhong, Q., et al., Edgetic perturbation models of human inherited disorders. Molecular systems biology, 2009. 5(1).

26. Madhani, H.D., C.A. Styles, and G.R. Fink, MAP kinases with distinct inhibitory functions impart signaling specificity during yeast differentiation. Cell, 1997. 91(5): p. 673–684.

27. Wang, X., et al., Three-dimensional reconstruction of protein networks provides insight into human genetic disease. Nature biotechnology, 2012. 30(2): p. 159.

28. Vidal, M., M.E. Cusick, and A.-L. Barabási, Interactome networks and human disease. Cell, 2011. 144(6): p. 986–998.

29. Landrum, M.J., et al., ClinVar: improvements to accessing data. Nucleic acids research, 2020. 48(D1): p. D835–D844.

30. Cui, H., N. Zhao, and D. Korkin, Multilayer View of Pathogenic SNVs in Human Interactome through In Silico Edgetic Profiling. Journal of molecular biology, 2018. 430(18): p. 2974–2992.

31. Subramanian, S., The abundance of deleterious polymorphisms in humans. Genetics, 2012. 190(4): p. 1579–1583.

32. Boyko, A.R., et al., Assessing the evolutionary impact of amino acid mutations in the human genome. PLoS genetics, 2008. 4(5): p. e1000083.

33. Tate, J.G., et al., COSMIC: the catalogue of somatic mutations in cancer. Nucleic acids research, 2019. 47(D1): p. D941–D947.

34. Dietlein, F., et al., Identification of cancer driver genes based on nucleotide context. Nature Genetics, 2020. 52(2): p. 208–218.

35. Hamosh, A., et al., Online Mendelian Inheritance in Man (OMIM), a knowledgebase of human genes and genetic disorders. Nucleic acids research, 2005. 33(suppl_1): p. D514–D517s.

36. Stenson, P.D., et al., Human gene mutation database (HGMD®): 2003 update. Human mutation, 2003. 21(6): p. 577–581.

37. Macgregor, S., et al., Associations of ADH and ALDH2 gene variation with self report alcohol reactions, consumption and dependence: an integrated analysis. Human molecular genetics, 2009. 18(3): p. 580–593.

38. Agarwal, D.P. and H.W. Goedde, Human aldehyde dehydrogenases: Their role in alcoholism. Alcohol, 1989. 6(6): p. 517–523.

39. Wall, T.L., et al., Hangover symptoms in Asian Americans with variations in the aldehyde dehydrogenase (ALDH2) gene. Journal of studies on alcohol, 2000. 61(1): p. 13–17.

40. Cook, T.A., et al., Associations of ALDH2 and ADH1B genotypes with response to alcohol in Asian Americans. Journal of Studies on Alcohol, 2005. 66(2): p. 196–204.

41. Eng, M.Y., S.E. Luczak, and T.L. Wall, ALDH2, ADH1B, and ADH1C genotypes in Asians: a literature review. Alcohol Research & Health, 2007. 30(1): p. 22.

42. Ye, L., Alcohol and the Asian flush reaction. SURG Journal, 2009. 2(2): p. 34–39.

43. Matoba, N., et al., GWAS of 165,084 Japanese individuals identified nine loci associated with dietary habits. Nature human behaviour, 2020. 4(3): p. 308–316.

44. Shankarkumar, U., The human leukocyte antigen (HLA) system. International Journal of Human Genetics, 2004. 4(2): p. 91–103.

45. Hildebrand, W.H., et al., HLA-B15: a widespread and diverse family of HLA-B alleles. Tissue antigens, 1994. 43(4): p. 209–218.

46. Bihl, F., et al., Impact of HLA-B alleles, epitope binding affinity, functional avidity, and viral coinfection on the immunodominance of virus-specific CTL responses. The Journal of Immunology, 2006. 176(7): p. 4094–4101.

47. Williams, F., et al., Analysis of the distribution of HLA-B alleles in populations from five continents. Human immunology, 2001. 62(6): p. 645–650.

48. Khan, M.A., HLA-B27 and its subtypes in world populations. Current opinion in rheumatology, 1995. 7(4): p. 263–269.

49. Goldberg, D.S. and F.P. Roth, Assessing experimentally derived interactions in a small world. Proceedings of the National Academy of Sciences, 2003. 100(8): p. 4372–4376.

50. Lo, C.-Y.Z., et al., Randomization and resilience of brain functional networks as systems-level endophenotypes of schizophrenia. Proceedings of the National Academy of Sciences, 2015. 112(29): p. 9123–9128.

51. Barabasi, A.-L. and Z.N. Oltvai, Network biology: understanding the cell’s functional organization. Nature reviews genetics, 2004. 5(2): p. 101.

52. Barabási, A.-L., N. Gulbahce, and J. Loscalzo, Network medicine: a network-based approach to human disease. Nature reviews genetics, 2011. 12(1): p. 56.

53. Choobdar, S., et al., Assessment of network module identification across complex diseases. bioRxiv, 2019: p. 265553.

54. Ghiassian, S.D., J. Menche, and A.-L. Barabási, A DIseAse MOdule Detection (DIAMOnD) algorithm derived from a systematic analysis of connectivity patterns of disease proteins in the human interactome. PLoS computational biology, 2015. 11(4): p. e1004120.

55. Will, C.L. and R. Lührmann, Spliceosome structure and function. Cold Spring Harbor perspectives in biology, 2011. 3(7): p. a003707.

56. Arzalluz-Luque, Á., et al., Mutant PRPF8 causes widespread splicing changes in spliceosome components in retinitis pigmentosa patient iPSC-derived RPE cells. Frontiers in Neuroscience, 2021. 15: p. 636969.

57. Kurtovic-Kozaric, A., et al., PRPF8 defects cause missplicing in myeloid malignancies. Leukemia, 2015. 29(1): p. 126–136.

58. Zhang, T., et al., SNRNP200 mutations cause autosomal dominant retinitis pigmentosa. Frontiers in Medicine, 2021. 7: p. 588991.

59. Zhang, X., et al., Contribution of SNRNP200 sequence variations to retinitis pigmentosa. Eye, 2013. 27(10): p. 1204–1213.

60. Kong, M.H., et al., Systematic review of the incidence of sudden cardiac death in the United States. Journal of the American College of Cardiology, 2011. 57(7): p. 794–801.

61. Offerhaus, J.A., C.R. Bezzina, and A.A. Wilde, Epidemiology of inherited arrhythmias. Nature Reviews Cardiology, 2019: p. 1–11.

62. Ruan, Y., N. Liu, and S.G. Priori, Sodium channel mutations and arrhythmias. Nature Reviews Cardiology, 2009. 6(5): p. 337.

63. Makita, N., et al., Novel calmodulin mutations associated with congenital arrhythmia susceptibility. Circulation: Cardiovascular Genetics, 2014. 7(4): p. 466–474.

64. Hennessey, J.A., et al., FGF12 is a candidate Brugada syndrome locus. Heart rhythm, 2013. 10(12): p. 1886–1894.

65. Musa, H., et al., SCN5A variant that blocks fibroblast growth factor homologous factor regulation causes human arrhythmia. Proceedings of the National Academy of Sciences, 2015. 112(40): p. 12528–12533.

66. Kataka, E., et al., Edgetic perturbation signatures represent known and novel cancer biomarkers. Scientific reports, 2020. 10(1): p. 1–16.

67. Mosca, R., et al., dSysMap: exploring the edgetic role of disease mutations. Nature methods, 2015. 12(3): p. 167–168.

68. Wang, K., M. Li, and H. Hakonarson, ANNOVAR: functional annotation of genetic variants from high-throughput sequencing data. Nucleic acids research, 2010. 38(16): p. e164–e164.

69. Das, J. and H. Yu, HINT: High-quality protein interactomes and their applications in understanding human disease. BMC systems biology, 2012. 6(1): p. 92.

70. Luck, K., et al., A reference map of the human protein interactome. bioRxiv, 2019: p. 605451.

71. Benedix, A., et al., Predicting free energy changes using structural ensembles. Nature methods, 2009. 6(1): p. 3–4.

72. Moal, I.H. and J. Fernández-Recio, SKEMPI: a Structural Kinetic and Energetic database of Mutant Protein Interactions and its use in empirical models. Bioinformatics, 2012. 28(20): p. 2600–2607.

73. Sussman, J.L., et al., Protein Data Bank (PDB): database of three-dimensional structural information of biological macromolecules. Acta Crystallographica Section D: Biological Crystallography, 1998. 54(6): p. 1078–1084.

74. Fiser, A. and A. Šali, Modeller: generation and refinement of homology-based protein structure models, in Methods in enzymology. 2003, Elsevier. p. 461–491.

75. Sondka, Z., et al., The COSMIC Cancer Gene Census: describing genetic dysfunction across all human cancers. Nature Reviews Cancer, 2018. 18(11): p. 696–705.

76. Eisenberg, E. and E.Y. Levanon, Human housekeeping genes, revisited. TRENDS in Genetics, 2013. 29(10): p. 569–574.

77. Zhu, J., et al., On the nature of human housekeeping genes. Trends in genetics, 2008. 24(10): p. 481–484.

78. Butte, A.J., V.J. Dzau, and S.B. Glueck, Further defining housekeeping, or “maintenance,” genes Focus on “A compendium of gene expression in normal human tissues”. Physiological genomics, 2001. 7(2): p. 95–96.

79. Kimura, M., Evolutionary rate at the molecular level. Nature, 1968. 217(5129): p. 624-626.

80. Kinsella, R.J., et al., Ensembl BioMarts: a hub for data retrieval across taxonomic space. Database, 2011. 2011.

81. UniProt: the universal protein knowledgebase. Nucleic acids research, 2017. 45(D1): p. D158–D169.

82. Ashburner, M., et al., Gene Ontology: tool for the unification of biology. Nature genetics, 2000. 25(1): p. 25–29.

83. Huang, D.W., B.T. Sherman, and R.A. Lempicki, Bioinformatics enrichment tools: paths toward the comprehensive functional analysis of large gene lists. Nucleic acids research, 2009. 37(1): p. 1–13.

84. Huang, D.W., B.T. Sherman, and R.A. Lempicki, Systematic and integrative analysis of large gene lists using DAVID bioinformatics resources. Nature protocols, 2009. 4(1): p. 44.

85. Freeman, L.C., A set of measures of centrality based on betweenness. Sociometry, 1977: p. 35–41.

86. Koschützki, D. and F. Schreiber, Centrality analysis methods for biological networks and their application to gene regulatory networks. Gene regulation and systems biology, 2008. 2: p. GRSB. S702.

87. Mizruchi, M.S., et al., Techniques for disaggregating centrality scores in social networks. Sociological methodology, 1986. 16: p. 26–48.

88. Puzis, R., et al., Augmented betweenness centrality for environmentally aware traffic monitoring in transportation networks. Journal of Intelligent Transportation Systems, 2013. 17(1): p. 91–105.

89. Girvan, M. and M.E. Newman, Community structure in social and biological networks. Proceedings of the national academy of sciences, 2002. 99(12): p. 7821–7826.

90. Newman, M.E., A measure of betweenness centrality based on random walks. Social networks, 2005. 27(1): p. 39–54.

91. Latora, V. and M. Marchiori, Efficient behavior of small-world networks. Physical review letters, 2001. 87(19): p. 198701.

92. Tripathi, S., et al., Comparison of module detection algorithms in protein networks and investigation of the biological meaning of predicted modules. BMC bioinformatics, 2016. 17(1): p. 129.

93. Vlaic, S., et al., ModuleDiscoverer: Identification of regulatory modules in protein-protein interaction networks. Scientific reports, 2018. 8(1): p. 433.

94. Cao, M., et al., Going the distance for protein function prediction: a new distance metric for protein interaction networks. PloS one, 2013. 8(10).

95. Cao, M., et al., New directions for diffusion-based network prediction of protein function: incorporating pathways with confidence. Bioinformatics, 2014. 30(12): p. i219–i227.

96. Pedregosa, F., et al., Scikit-learn: Machine learning in Python. the Journal of machine Learning research, 2011. 12: p. 2825–2830.

