## Supplementary Information for "The Extent of Edgetic Perturbations in the Human Interactome Caused by Population-Specific Mutations"

### Edgotype Analysis of SNVs Reveals Variance across the Human Interactome Landscape and Population Phenotypes

### Supplementary Information

#### List of figures:


| | |
|---|---|
| **Supplementary Figure S1** | Top 20 enriched GO terms in three basic GO categories: biological process, molecular function and cellular component |
| **Supplementary Figure S2** | HLA-A is one of genes carrying high number of normal disruptive mutation in the HLA gene family |
| **Supplementary Figure S3** | Supplementary Figure S3. Distinct rewiring patterns in interactome modules associated with arrhythmias in East Asians and Americans. |
| **Supplementary Figure S4** | Illustration of the concept of Diffusion State Distance (DSD) |


#### List of tables:


| | |
|---|---|
| **Supplementary Table S1** | Disease phenotype information associated with genes enriched with disruptive mutation curated from OMIM, HGMD |

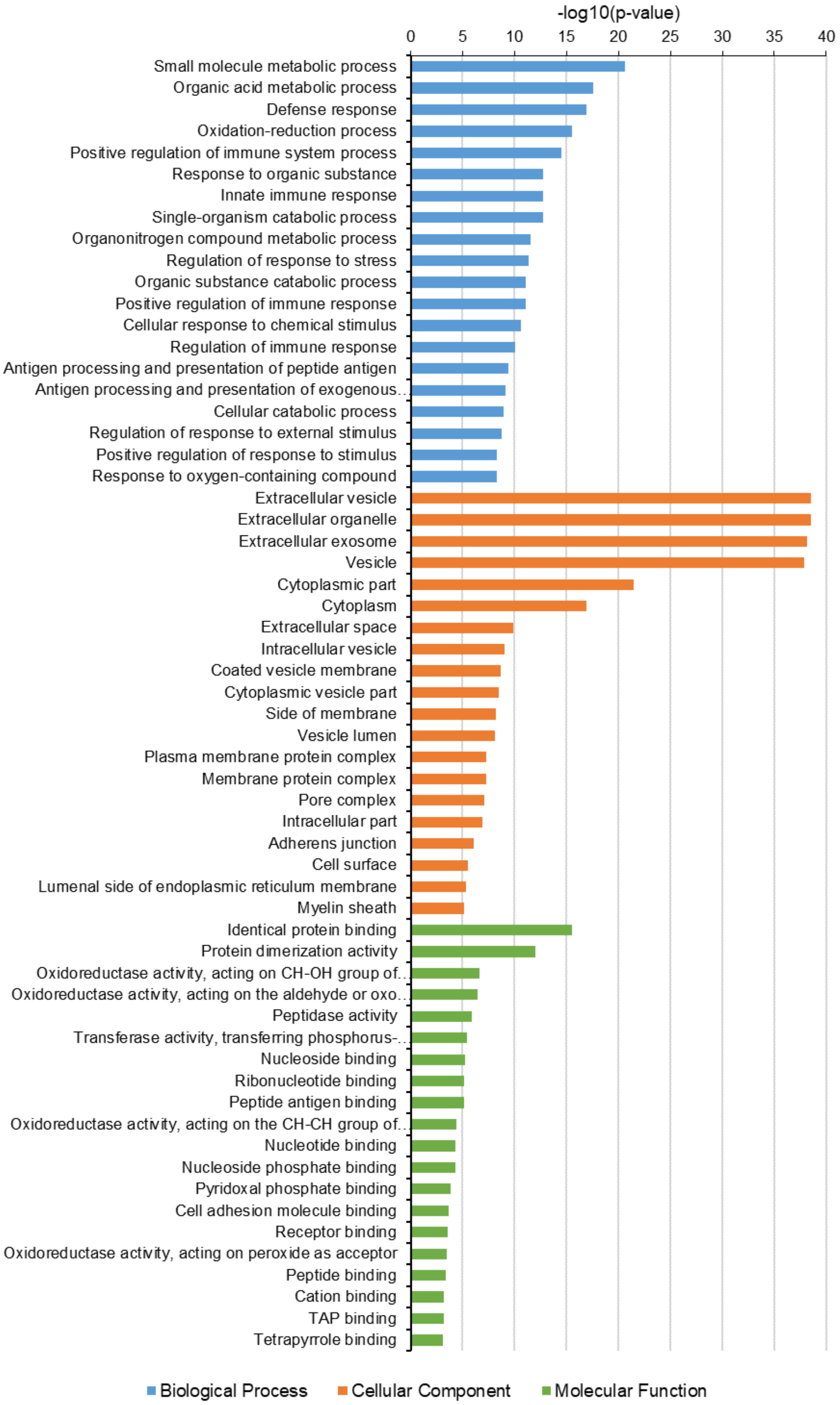


**Supplementary Figure S1. Top 20 significant GO terms for genes enriched with disruptive mutations among three basic GO categories: biological process, molecular function and cellular component.**

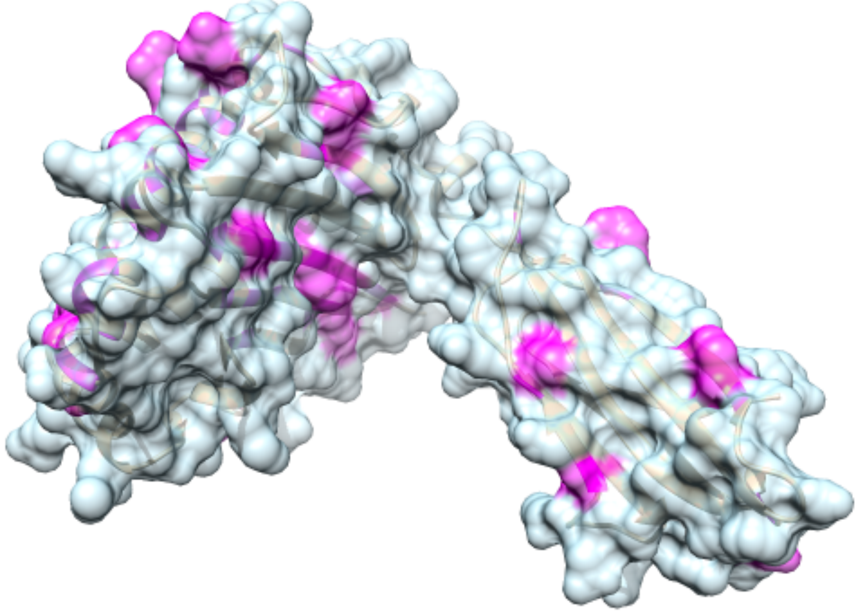

**Supplementary Figure S2. HLA-A is one of the top genes carrying high number of normal disruptive mutation in the HLA gene family.**

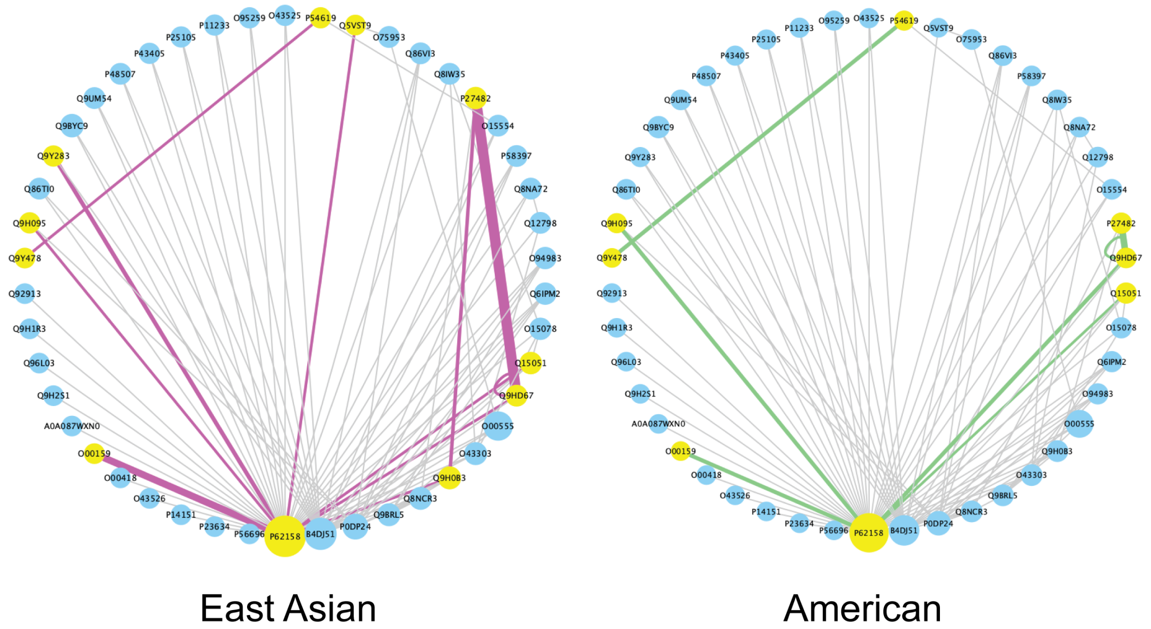


**Supplementary Figure S3. Distinct rewiring patterns in interactome modules associated with arrhythmias in East Asians and Americans.** The East Asian population has 11 proteins that carry mutations (yellow nodes) resulting in 11 rewired interactions (magenta edges), whereas the American population has 6 mutant proteins (yellow nodes) with 6 rewired interactions (green edges). Additionally, the higher allele frequency of a mutation and the corresponding rewired interaction is represented by the increasing thickness of edges, where one can see that the East Asian population has higher frequency of these mutations.

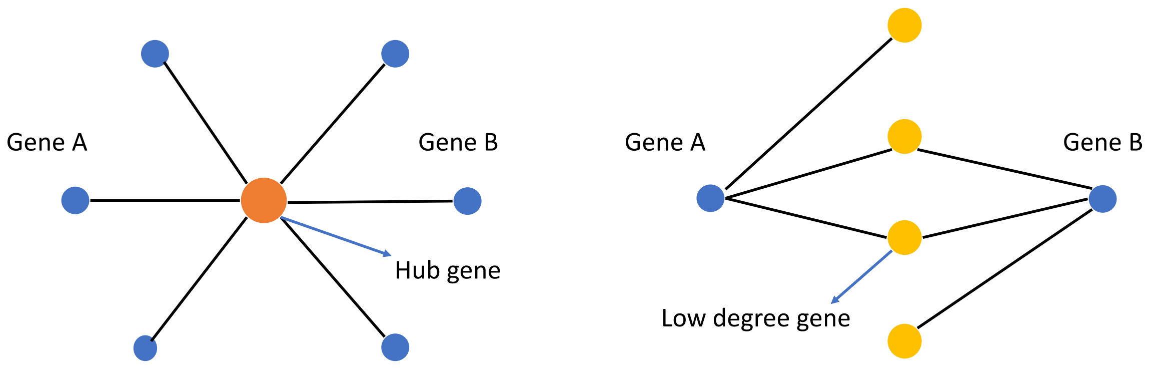


**Supplementary Figure S4 Illustration of the concept of Diffusion State Distance (DSD).** DSD downweighs the influence of the hub gene and favors low degree gene.
